# Folding PDZ2 domain using the Molecular Transfer Model

**DOI:** 10.1101/037283

**Authors:** Zhenxing Liu, Govardhan Reddy, D. Thirumalai

## Abstract

A major challenge in molecular simulations is to describe denaturant-dependent folding of proteins order to make direct comparisons with *in vitro* experiments. We use the molecular transfer model (MTM), which is currently the only method that accomplishes this goal albeit phenomenologically, to quantitatively describe urea-dependent folding of PDZ domain, which plays a significant role in molecular recognition and signaling. Experiments show that urea-dependent unfolding rates of the PDZ2 domain exhibit a downward curvature at high urea concentrations ([*C*]s), which has been interpreted by invoking the presence of a sparsely populated high energy intermediate. Simulations using the MTM and a coarse-grained Self-Organized Polymer (SOP) representation of PDZ2 are used to show that the intermediate (*I_EQ_*), which has some native-like character, is present in equilibrium both in the presence and absence of urea. The free energy profiles as a function of the structural overlap order parameter show that there are two barriers separating the folded and unfolded states. Structures of the transition state ensembles, (*TSE*1 separating the unfolded and *I_EQ_* and *TSE*2 separating *I_EQ_* and the native state), determined using the *P_fold_* method, show that *TSE*1 is greatly expanded while *TSE*2 is compact and native-like. Folding trajectories reveal that PDZ2 folds by parallel routes. In one pathway folding occurs exclusively through *I*_1_, which resembles *I_EQ_*. In a fraction of trajectories, constituting the second pathway, folding occurs through a combination of *I_1_* and a kinetic intermediate. We establish that the radius of gyration (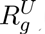) of the unfolded state is more compact (by ∼ 9%) under native conditions. Theory and simulations show that the decrease in 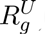 occurs on the time scale on the order of utmost ~ 20 *μβ.* The modest decrease in 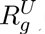 and the rapid collapse suggest that high spatial and temporal resolution, currently beyond the scope of most small angle X-ray scattering experiments, are needed to detect compaction in finite-sized proteins. The present work further establishes that MTM is efficacious in producing nearly quantitative predictions for folding of proteins under conditions used to carry out experiments.

## Introduction

The Molecular Transfer Model (MTM)^1^, based on the statistical mechanical theory of liquid mixtures^2^, is currently the only available computational method that predicts the outcomes of experiments and provides the structural basis of folding as a function of denaturants^3,4^ and pH^5^. Using the MTM in conjunction with coarse-grained (CG) representation of polypeptide chains we have made quantitative predictions of the folding thermodynamics and kinetics as a function of denaturants for a number of small (protein L, cold shock protein, srcSH3 domain, and Ubiquitin)^1,2,4,6^, and GFP, a large single domain proteins^3^. Because the effects of denaturants are taken into account naturally within the MTM framework^2^, albeit phenomenologically it has been possible to obtain chevron plots for src SH3 domain producing quantitative agreement with experiments for the slopes of folding and unfolding arms^7^. Although MTM can be be implemented in conjunction with atomically detailed simulations we have so far used CG models for proteins. The virtue of CG models^8-11^ is that they can be used to obtain both the thermodynamic and kinetic properties over a a wide range of external conditions, thus allowing us to compare with experiments directly^12^. These studies illustrate that simulations based on the MTM provide concrete predictions for *in vitro* experiments enabling us to go beyond generic ideas used to understand protein folding^13-23^. Here, we investigate the folding mechanism of PDZ2 domain using CG simulations within the theoretical framework of the MTM.

PDZ domains are a large family of globular proteins that mediate protein-protein interactions and play an important role in molecular recognition^24-26^. These proteins generally consist of 80 —100 amino acids. PDZ2 domain has six *β* strands and two *α* helices (Figure 1a). The folding mechanism of PDZ2, with 94 residues, has been studied experimentally^27^ using classical chemical kinetics methods. The key findings in these experiments are: (i) In urea-induced equilibrium denaturation experiments, the observed transition is cooperative, which is well described by an apparent two-state model^27,28^. (ii) In a majority of cases proteins that fold thermodynamically in a two-state manner also exhibit a similar behavior kinetically in ensemble experiments. However, urea-dependent unfolding rates exhibit a downward curvature at high urea concentrations at pH > 5.5. Based on the observation that the folding kinetics is mono-phasic, with no detectable burst phase in the initial fluorescence of the initial unfolding time course, it was surmised that there is no low energy intermediate in the unfolding of PDZ2. Rather the data were used to suggest the presence of a high-energy on-pathway intermediate, which does not accumulate significantly in equilibrium^27^. (iii) The high energy intermediate is non-detectable under stabilizing conditions, achievable in PDZ2 domain by addition of modest amount of sodium sulfate. Under these conditions PDZ2 folds thermodynamically and kinetically in a two-state manner. (iv) The structures of the two transition state ensembles were also inferred using measured Φ values as constraints in all atom molecular dynamics simulations^29^. Unlike the results summarized in (i)- (iii) the predicted structures of the transition state ensembles are not as conclusive for reasons explained later in this work.

**Figure 1:**
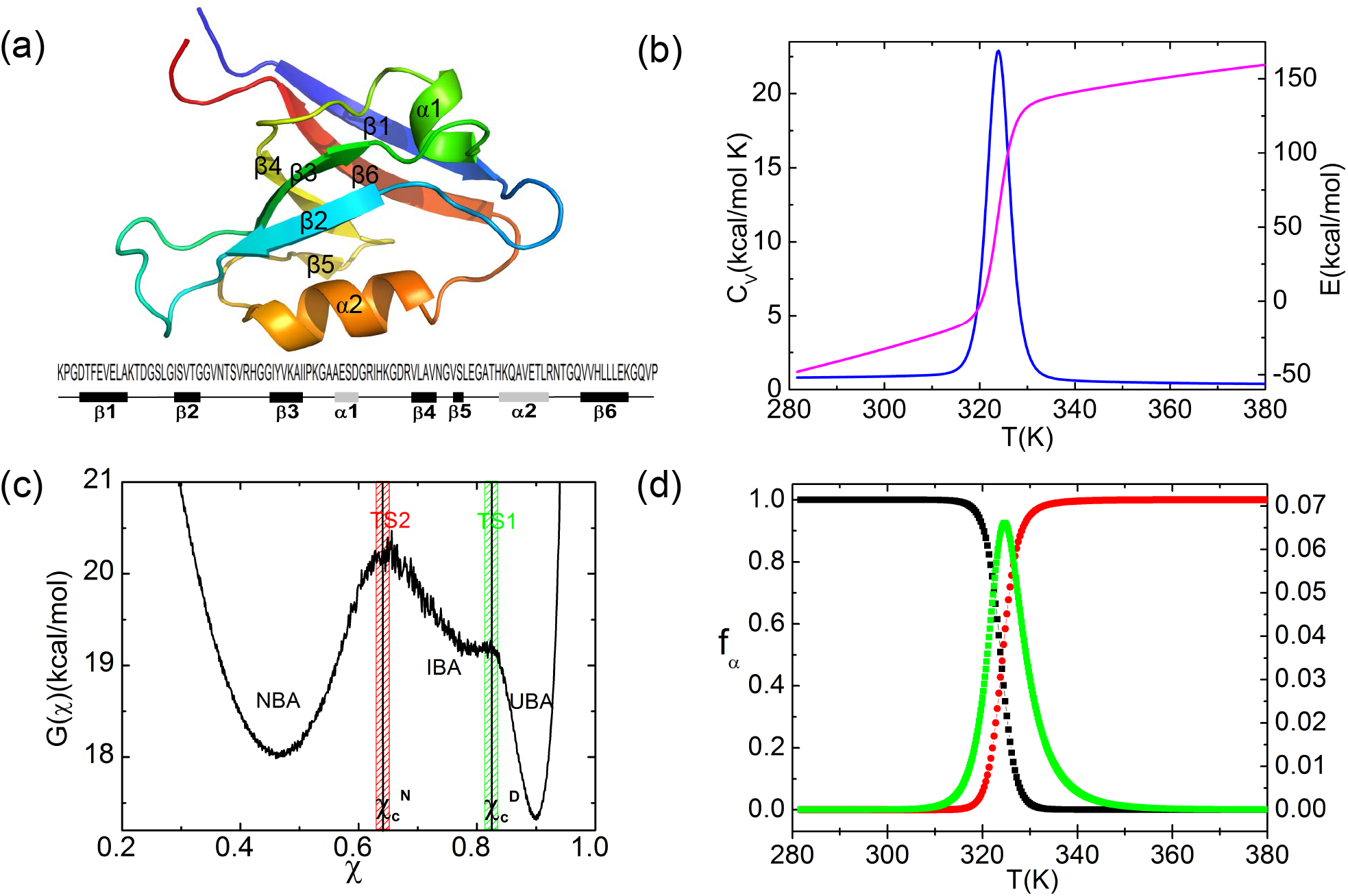
Thermodynamics of folding. (a) Ribbon diagram representation of PDZ2(PDB code: 1GM1). (b) Temperature dependence of specific heat (blue) and total energy (magenta). (c) Free energy profile at *T_m_* as a function of *χ*. The values 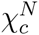 and 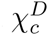 are used to classify the major equilibrium states. The shaded areas give putative regions for the two transition state ensembles. (d) Fraction of molecules in NBA(black), UBA(red) and IBA(green) as functions of temperature.

In order to provide a comprehensive picture of folding of PDZ2, with potential implications for other single domain proteins, we performed molecular simulations of a coarse-grained off-lattice model with side chains^9,30^ and used MTM^1,3,7^ to account for for denaturant effects. The free energy profiles as a function of the structural overlap order parameter at different urea concentrations, [*C*], and temperature *T* suggest that the folding mechanism of PDZ2 can be altered by changing the stability of the folded state. In accord with experiments, we demonstrate directly the existence of the fleeting obligatory intermediate both in equilibrium (*I_EQ_*) and kinetics (*I*_1_)^31^. The structures of *I_EQ_* and *I*_1_ are similar. However, the fraction of molecules in intermediate basin of attraction (IBA),*f_IBA_*, as a function of temperature and [*C*] is small, thus explaining the difficulty in detecting it in standard denaturation experiments. In addition to *I*_1_ a kinetic intermediate, *I*_2_, is consistently populated in ∼ 53% of the folding trajectories. Guided by the free energy profiles, we identified two transition state ensembles, *TSE*1 and *TSE*2. The computed values of the Tanford-like *β* parameters, using the solvent accessible surface area as a surrogate, for the two transition state ensembles (one connecting the NBA and the IBA and the other involving transition between the IBA and the UBA) are in qualitative agreement with those obtained from experiments^27,31^. The current work further establishes that simulations based on the MTM are efficacious in providing a nearly quantitative picture of folding of single domain proteins.

## METHODS

**SOP-sidechain model**: The simulations were carried out using a CG model in which the *C_α_*-based self-organized polymer (SOP) representation^32^ was augmented to include side chains(SCs)^2,7^. In the SOP-SC model each residue is represented by two interaction centers, one that is located at the *C_α_* position and the other at the center of mass of the side chain. In the SOP-SC model the native state stabilization is achieved by accounting for backbone-backbone (bb), side chain-side chain (ss), and backbone-side chain (bs) interactions present in the folded state. Neglect of non-native interactions, which do not significantly alter the folding mechanism beyond the global collapse of the protein^33-36^, is nevertheless a limitation of the model.

The energy (to be interpreted as an effective free energy obtained by integrating over solvent (water) degrees of freedom) of a conformation, describing the intra peptide interactions, is

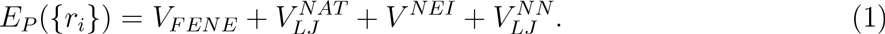

The finite extensible nonlinear elastic potential (FENE), *V_FENE_*, accounting for the chain connectivity between backbones and side chains, is,

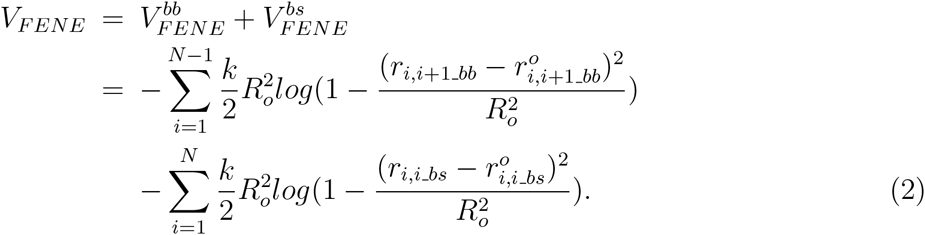

The non-bonded native interaction, 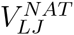 in Eq. (1) is taken to be

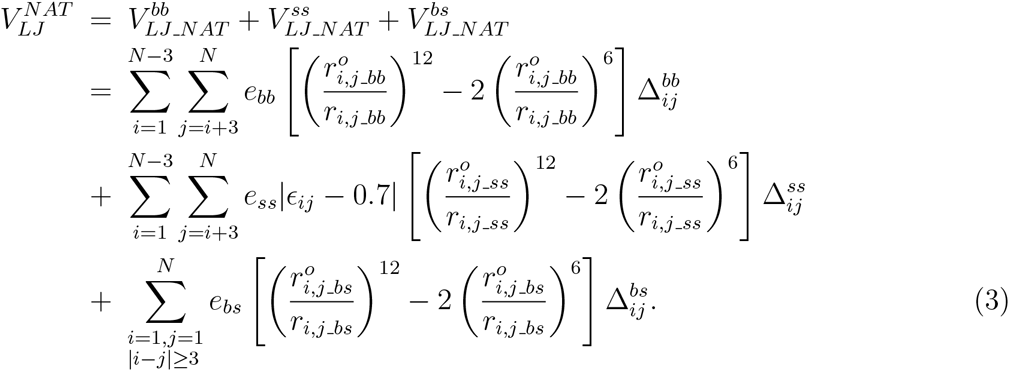

In Eqs. 2 and 3 the superscript *^o^* refers to distances in the native state. If the distance between two non-covalently linked beads, *r_ij_*(|*i* — *j*| ≥ 3) in the PDB structure is within a cutoff distance *R_c_*, a native contact is formed, and correspondingly *Δ_ij_* = 1. If *r_ij_* exceeds *R_c_* then *Δ_ij_* = 0. The strengths of the non-bonded interactions *e_bb_*, *e_ss_*, *e_bs_* are assumed to be uniform. The Betancourt-Thirumalai (BT)^37^ statistical potential matrix with elements *є_ij_*, is used to explicitly treat the sequence dependence.

We used repulsive interactions for excluded volume effects between neighboring beads with strength *e_l_*. The ranges of repulsion are *σ_bb_*, *σ_i,j_ss_*, *σ_j_bs_* for bb, ss and bs interactions respectively. The form of *V^NEI^* is

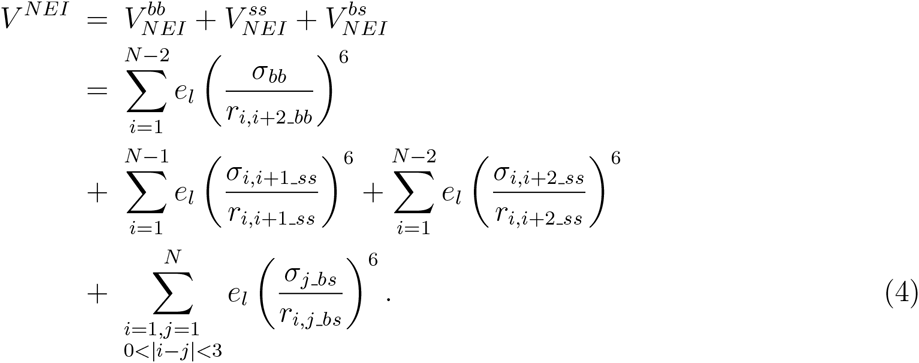

To prevent interchain crossing, we choose *σ_bb_* = *a* = 3.8*Å* (a is average distance between neighboring *C_α_* atoms), *σ_ij_ss_*, = *f(σ_i_* + *σ_j_*) *(σ_i_*, *σ_j_* are the van der Waals radii of the side chains and *f* = 0.5), *σ_j_bs_* = *f(a + σj*).

The non-bonded nonnative interactions are given by

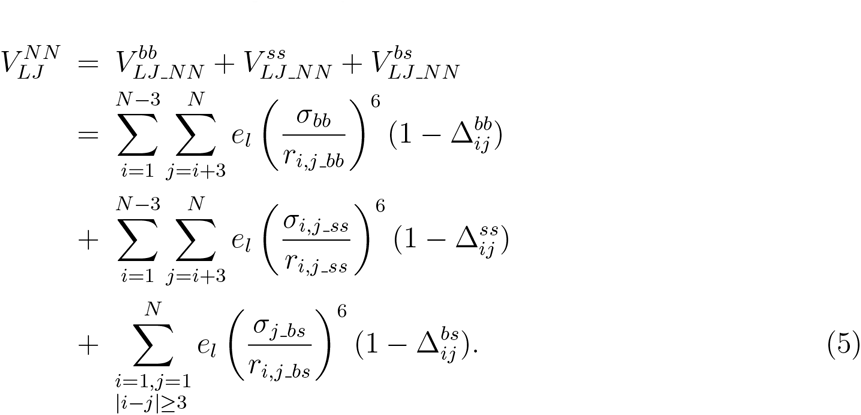

Besides the knowledge-based BT statistical potential, the SOP-SC energy function *E_P_({r_i_}*) has seven parameters: *R_o_* = 2*Å*, *k* = *20kcal/(mol · Å*^2^), *R_c_* = 8*Å*, *e_bb_* = *0.73kcal/mol, e_bs_* = *0.17kcal/mol, e_ss_* = *0.3kcal/mol, e_i_* = *lkcal/mol.* Among these parameters, *R_o_* and *k* merely account for chain connectivity. The results would not be significantly affected by the precise choice of parameters enforcing the integrity of the polypeptide chain. Thus, in effect there are five parameters in the SOP-SC model. Analysis of PDB structures shows that *R_c_* = 8*Å* is a reasonable choice. By analyzing structures of folded proteins in the Protein Data Bank (PDB) that there are large favorable bb and bs contacts^38^. The first and third terms account for these interactions, which in turn ensures that packing effects are appropriately described in the SOP-SC model. The experimental melting temperature of the protein is used to determine the strengths of the native contacts, specified by *e_bb_, e_bs_, e_ss_.*

***Molecular Transfer Model***: The MTM theory^2^ shows that experimentally measured transfer free energies for backbone and side chains along with the SOP-SC model could be utilized to obtain the partition function for a protein at a finite denaturant concentration [C]^2^. In the MTM, the free energy of transferring a given protein conformation, described by {*r_i_*} from water ([C]=0) to aqueous denaturant solution ([C]≠0), is approximated as,

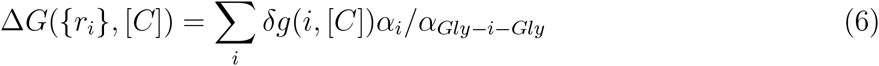

where the sum is over backbone and side chain, *δg(i, [C])* is the experimentally measured transfer free energy of group *i*, *α_i_* is the solvent accessible surface area (SASA), and *α_Gly-i-Gly_* is the SASA of the *i^th^* group in the tripeptide *Gly — i — Gly.* Thus, the effective free energy function for a protein at [C]=0 is

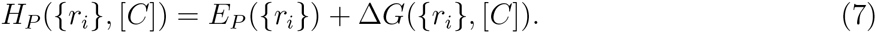

For computational expediency we combined converged simulations at finite temperature using *E_P_({r_i_})* and computed the partition function at [C]≠0 to obtain thermodynamic properties using a weighted histogram analysis method, which takes into account the effects of *ΔG({r_i_}, [C])*. Such an approximation, whose validity for obtaining thermodynamic properties has been previously established^7^, is used to obtain thermodynamic properties.

**Langevin and Brownian Dynamics Simulations**: We assume that the dynamics of the protein is governed by the Langevin equation, which includes a damping term with a friction coefficient ζ, and a Gaussian random force Γ. The equation of motion for a generalized coordinate *r_i_* is 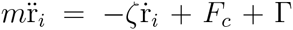 where *m* is the mass of a bead, *F_c_* = − *∂E_P_({r_i_})/∂r_i_*, is the conformational force calculated using Eq. (1), Γ is the random force with a white noise spectrum. The autocorrelation function for Γ(*t*) in the discretized form is 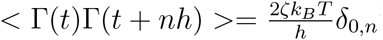^39^ where *δ_0_,_n_* is the Kronecker delta function and *n* = 0,1,2.… The value of the time step, *h*, depends on the friction coefficient *ζ*.

To obtain enhanced sampling, we used the Replica Exchange Molecular Dynamics (REMD)^40-43^ to perform thermodynamics sampling using a low friction coefficient ζ = 0.05m/T_L_^44^, which allows us to accurately calculate the equilibrium properties. Rapid convergence of thermodynamics is possible at a small ζ (under damped limit) because the polypeptide chain makes frequent transitions between all accessible states. In the low ζ limit limit, we used the Verlet leap-frog algorithm to integrate the equations of motion. The velocity at time t + *h/2* and the position at time *t* + *h* of a bead are given by,

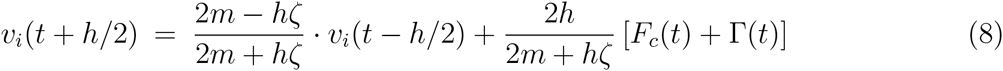

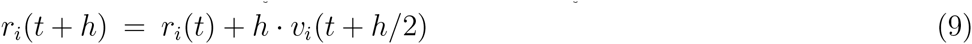

In order to simulate the kinetics of folding, ζ is set to be 50m/T_L_, which approximately corresponds to the value in water and represents the over damped limit^39^. In the high ζ value, we use the Brownian dynamics algorithm^45^, which allows us to integrate equations of motion using

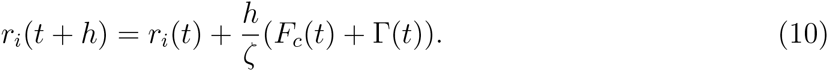

**Time Scales**: The natural unit of time for over damped condition at the transition temperature *T_s_* is 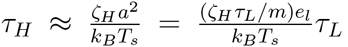. To convert the simulation time to real time, we chose *e_l_* = *lkcal/mol,* average mass *m* =1.8 x 10^-22^g^39^, *a* = 4*Å*, which makes *τ_L_ = 2ps.* For *ζ_Η_ = 50m/τ_L_*, we obtain *τ_H_ = 159ps*. For thermal folding simulations, the integration time step, *h* is 0.005*τ_L_*. In the kinetic folding simulations, *h*, in Eq.(10) is 0.02*τ_H_*.

***Data Analysis***: The [C]-dependent melting temperature is identified with the peak in the heat capacity. The structural overlap function 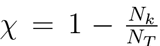 is used to monitor the folding reaction, where

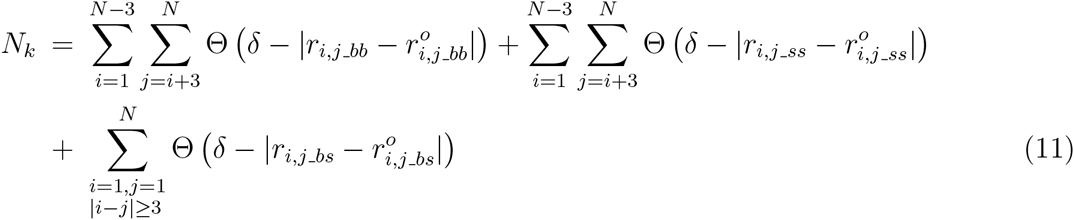

In Eq.(11), Θ(*x*) is the Heavyside function. If 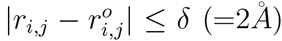, there is a contact. The number of contacts in the *k^th^* conformation is *N_k_*, and *N_T_* is the total number in the folded state.

## Results

**Thermal Denaturation:**: The temperature dependence of the heat capacity, *C_v_*(= 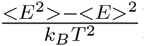, where < *E* > and < *E*^2^ > are the mean and mean square averages of the energy, respectively, demonstrates that PDZ2 folds cooperatively in a two-state manner (Figure 1b). The melting temperature, identified with the peak in *C_v_(T*), is *T_m_* = 324K. The value of *T_m_* obtained in our simulations is in excellent agreement with the experimentally measured *T_m_* = 321K^47^.

In order to identify the *NBA, UBA,* and the intermediate basin of attraction *(IBA)* representing the *I_EQ_* state, we plot in Figure 1c the free energy (G(χ)) profile as function of χ at *T_m_* = 324K. All the conformations can be classified into three states specified by the black vertical lines based on the χ values. If 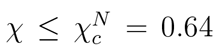, the conformations are in the NBA, conformations with 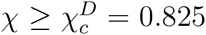 belong to the UBA, and the rest of the conformations are in the IBA. The fractions of molecules in the *NBA, f_NBA_([0],T)* (the first argument shows the value of the denaturant concentration), in the *UBA f_UBA_([0],T)*, and in the *IBA, f_1BA_([0],T)* as a function of temperature are plotted in Figure 1d. Both *f_NBA_([0],T*) and *f_UBA_([0],T)* show that the folding or unfolding is cooperative. The value of *f_IBA_([0],T)* is negligible compared to *f_NBA_([0],T)* and *f_UBA_([0],T)*, suggesting that from a thermodynamic perspective a two-state description is adequate, reflecting the cooperative transition in the heat capacity curve (Figure 1b). The sparse population of the IBA explains why the *I_EQ_* state is hard to detect in experiments although its presence appears as a shoulder in the free energy profile (Figure 1c). The value of *T_m_* computed using *f_NBA_(T_m_)* = 0.5 yields *T_m_* = 324K, which coincides with the peak in the heat capacity (Figure 1b).

**Chemical Denaturation:** Following our previous studies^1,3,7^, we choose a simulation temperature, *T_s_,* at which the calculated free energy difference between the native state (N) and the unfolded state (U), Δ*G_NU_(T_s_) *(G*_N_(T_s_) — G_U_(T_s_))* and the measured free energy Δ*G_NU_(T_E_)* at *T_E_* (=298K) coincide. The use of Δ*G_NU_(T_s_) = ΔG_NU_(T_E_)* ( in water with [*C*] = 0) to fix *T_s_* is equivalent to choosing a overall reference energy scale in the simulations. For PDZ2, Δ*G_NU_(TE = 298K) = —3.1kcal/mol* at [*C*] = 0^28^, which results in *T_s_ = 317K*. Besides the choice of *T_s_*, *no other parameter* is adjusted to obtain agreement with experiments for any property.

With *T_s_ = 317K* fixed, we calculated the dependence of *f_NBA_([C],T_s_), f_UBA_([C],T_s_)*, and *f_IBA_([C],T_s_)* on [*C*] (Figure 2a). The agreement between the measured and simulated results for *f_NBA_([C],T_s_)* as a function of [*C*] is excellent (Figure 2a). We find, just as in thermal denaturation, that *f_IBA_([C],T_s_)* is small^27,28^. The midpoint concentration, *C_m_*, obtained using *f_NBA_([C_m_],T_s_)* = 0.5 is [C]=2.3M, agrees well with the experimentally measured value of 2.6M (see Figure 6D in ^28^).

**Figure 2:**
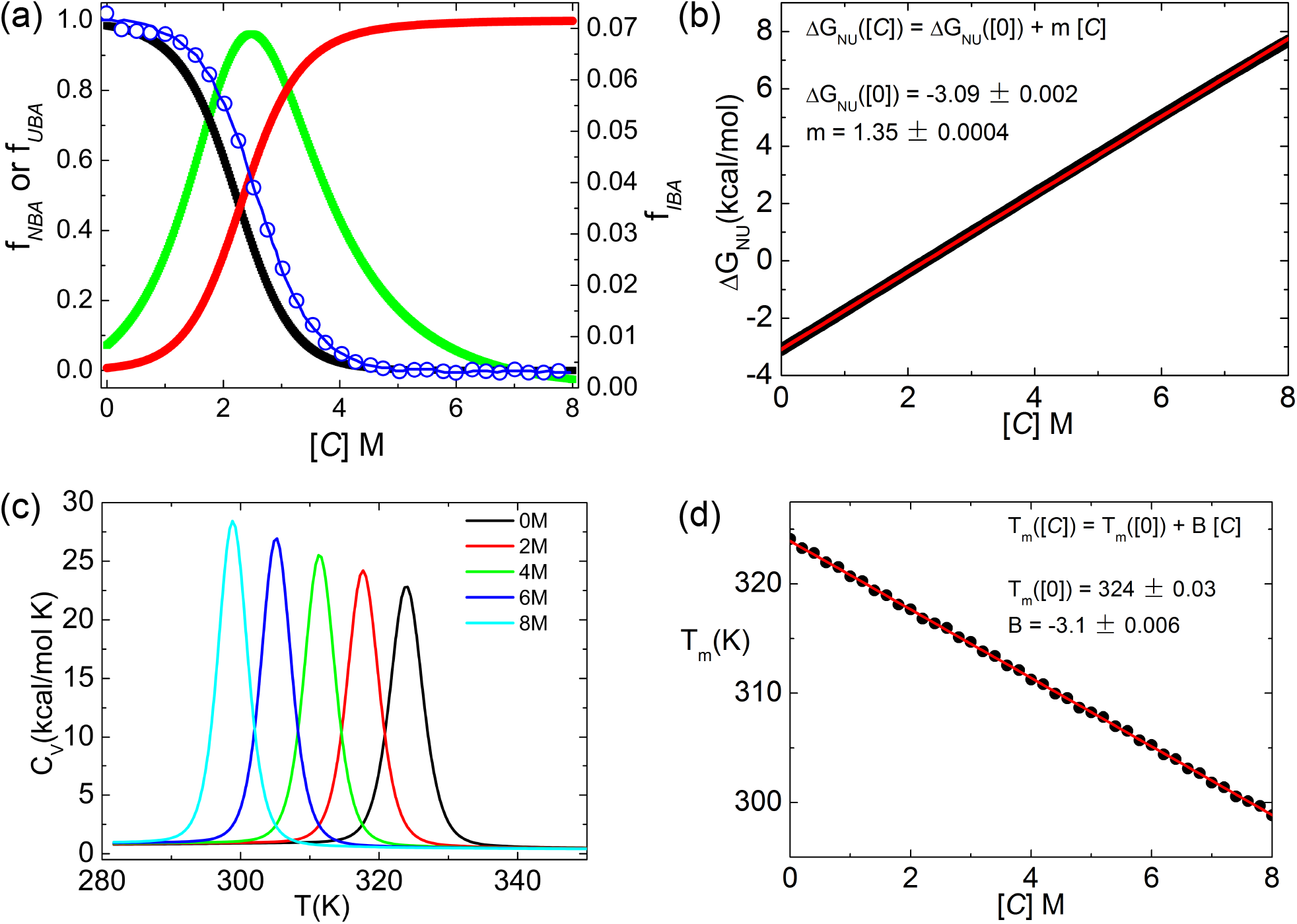
Denaturation effects. (a) Fraction of molecules in the NBA (black), UBA (red), and IBA (green) as a function of urea concentration [*C*]. For comparison, the experimental curve for *f_NBA_[C*] (blue) is shown. (b) [*C*] dependence of free energy of stability of the native state with respect to the unfolded state. Fit to a linear function yields Δ*G_NU_ = ΔG_NU_ (0) + m[C]* where Δ*G_NU_(0) = – 3.09kcal/mol* and *m* = *1.35kcal/mol.M*. (c) Heat capacity versus temperature for different values of [*C*]. (d) The [*C*] dependence of the melting temperature. The line is a fit to *T_m_[C*] = *T_m_*(0) — *B[C*] where *T_m_(0) = 324K* and *B* = *— 3.1KM*^-1^.

The native state stability with respect to U, Δ*G_NU_([C])(= G_N_([C]) — G_U_([C]))*, is computed using 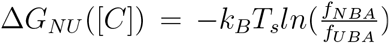. The linear fit, Δ*G_NU_([C]) = ΔG_NU_(0) + m[C]*, yields Δ*G_NU_([O]) = — 3.09kcal/mol* and *m = 1.35kcal/mol · M* (Figure 2b). The experimentally inferred *m = 1.20kcal/mol · M* compares well with the simulations, which establishes again that simulations based on MTM predict the thermodynamic properties of proteins accurately. The [*C*]-dependent heat capacity curves (Figure 2c) show that the peaks corresponding to *T_m_([C*]) decreases as [*C*] increases (Figure 2c=d). Taken together the simulations show that the equilibrium folding induced by temperature or denaturants is cooperative.

**Urea-dependent changes in the shape of PDZ2:** The dependence of the radius of gyration, 〈*R_g_*([*C*])〉, on urea concentration (black line in Figure 3a) shows a transition from an expanded to a collapsed state as [*C*] decreases. The radii of gyration of the *I_E_Q* and the native state are virtually constant at all urea concentrations. However, the radius of gyration of the *UBA* decreases as [*C*] decreases implying that the ensemble of structures of the unfolded state is more compact under native conditions than at 8M urea. The decrease in the *R_g_* of the *UBA* structures in going from 8M urea to 1M is about 9%, which should be measurable in a high precision Small Angle X-ray scattering (SAXS) experiment (see below for additional discussion). The *P(R*_g_) distributions at various urea concentrations show the expected behavior (Figure 3b). The protein is largely in the *NBA* at [*C*] less than 2.3M (mid point of the folding transition) and is expanded at higher values of [*C*]. The distance distributions, which can be measured as the inverse Fourier transform of the wave vector dependent scattering intensity, are plotted in Figure 3c and constitutes one of the predictions.

**Figure 3:**
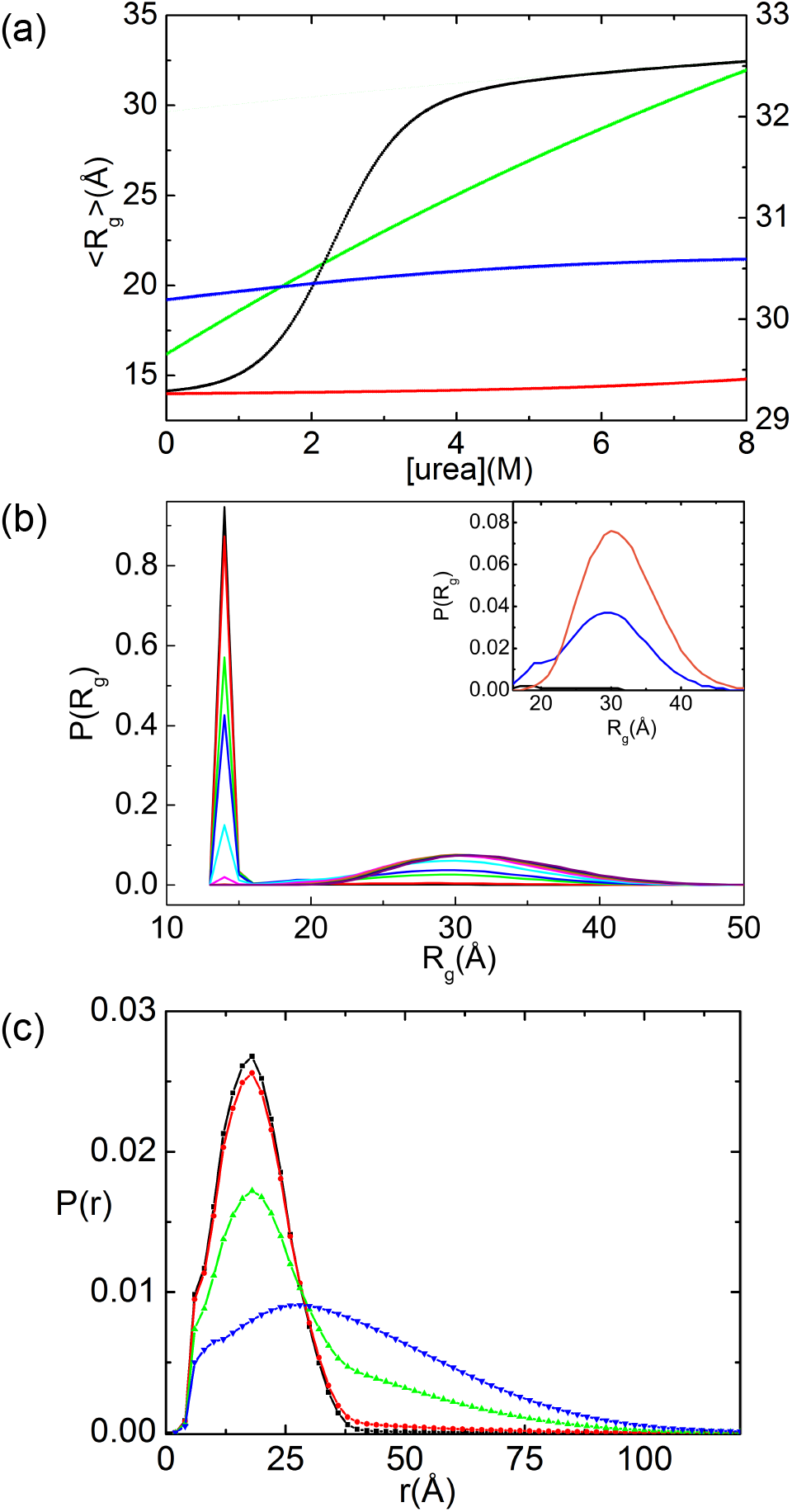
Equilibrium collapse. (a) Average < *R_g_* > (black) as a function of [*C*]. Red, green and blue curves correspond to < *R_g_* > of the folded, unfolded and *I_EQ_* states, respectively. The scale for the unfolded state is on the right. (b) Distribution *P(R_g_)* of *R_g_* for various concentrations of urea. The inset shows *P(R_g_)* for [*C*] = 0(black), [*C*] = 2.3(blue), and [*C*] = 5.0(orange) corresponding to the extended conformations *(R_g_* > 16Â). (c) Distance distribution function *P(r)*, the inverse Fourier transform of the scattering intensity, for 0M (black), 1.0M (red), 2.3M (green), and 5.0M (blue) urea. Here, *r* is the distance between all non-covalently linked beads.

**Free energy profiles,** *G(χ)*, **reveal lowly populated intermediate:** To illustrate how urea changes the folding landscape, we plotted *G(χ)* versus χ at different [*C*] at *T* = *T_m_* = 324K in Figure 4a and at *T* = *T_s_* = 317K in Figure 4b. Figure 4a shows, that at all urea concentrations, the conformations could be partitioned into three states, which is consistent with the results in Figure 1c displaying *G(χ)* at *T_m_* with [*C*] = 0. The basin of attraction corresponding to *I_EQ_* becomes deeper as *[C*] increases but is shallow compared to the *NBA* and *UBA*. The barrier for *TSE*1 is lower than for *TSE*2, implying that the transition from the intermediate states to the UBA is more facile than transition to the *NBA*. The number of distinct minima remain unchanged at the lower temperature, *T_s_* = 317K (Figure 4b). However, the basin for the intermediate states becomes shallower as [*C*] increases, and the barrier for TSE1 almost disappears, especially when [*C*] > *C_m_* = 2.3*M*, which indicates that the intermediate, if it can be detected at all, is unstable. The absence of *I_EQ_* at the lower temperature suggests that stabilization of the native fold of PDZ2 leads to a two-state thermodynamic transition. Our finding supports the same observation in experiments showing that native state stabilization upon addition of sodium sulfate results in the high energy *I_EQ_* being undetectable.

**Figure 4:**
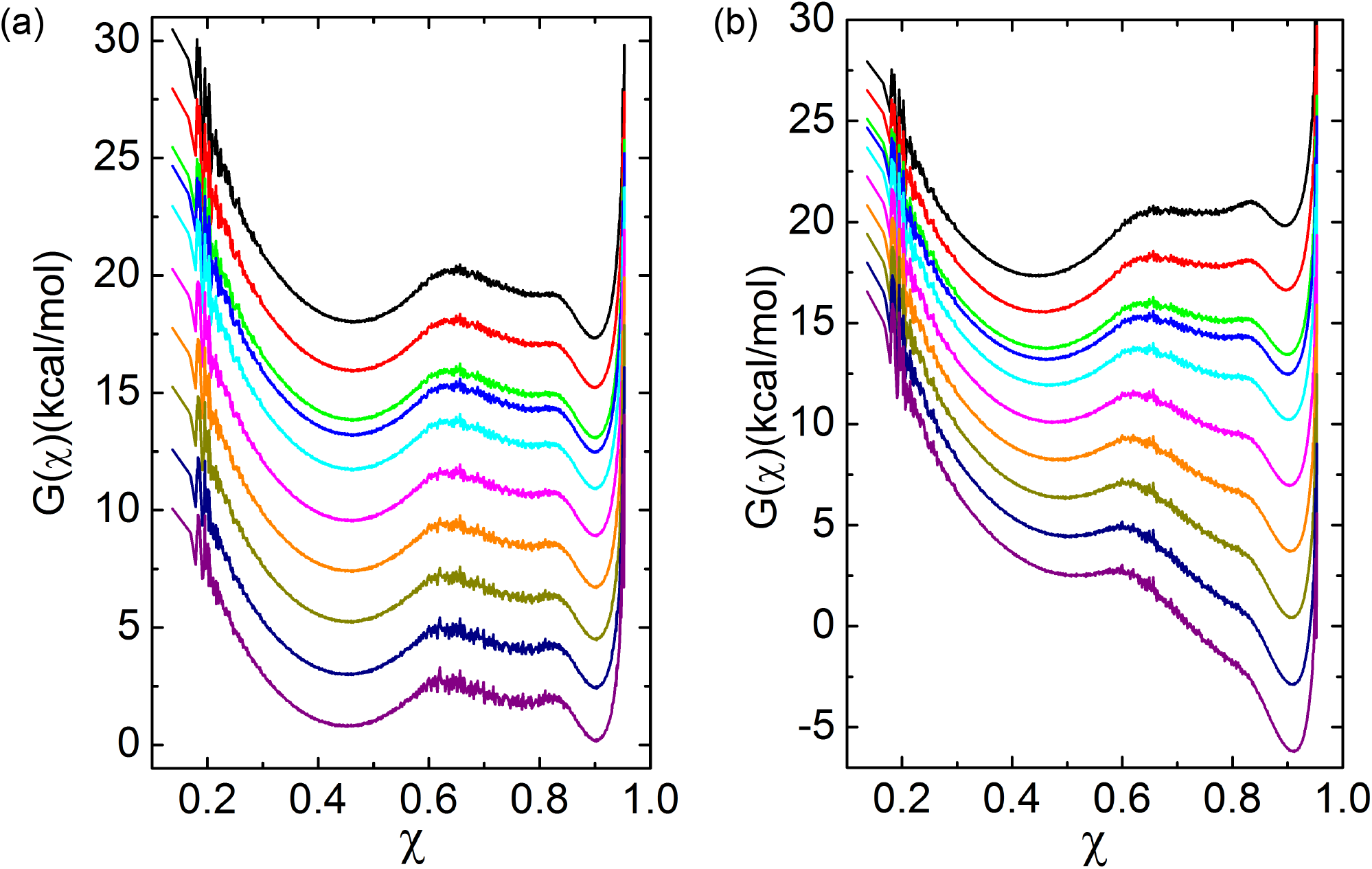
Free energy profiles versus *χ* at different [*C*]. (a) *T* = *T_m_.* (b) *T* = *T_s_.* The values of [*C*] measured in M from top to bottom are 0, 1, 2, 2.3, 3, 4, 5, 6, 7, and 8.

**Structures of the transition state ensembles** (*TSE*1 **and** *TSE*2): We identified the transition state structures using the folding trajectories generated at *T_m_*. We picked the putative transition state from the barrier regions of the free energy profile as a function of χ, using the conditions 0.815 < χ < 0.835 for *TSE*1 and 0.63 < χ < 0.65 for *TSE*2, which are represented as shaded areas in Figure 1c. Starting from these structures, we calculated the commitment probability, *P_fold_*, of reaching the *NBA^48^*. The sets of structures with 0.4 ≼ *P_fold_* ≼ 0.6 are identified as the transition state ensemble. The characteristics of the two transition state ensembles are displayed in Figure 5a and the distribution of the structural order parameter *P*(χ) of the *TSE*1 along with the contact map are shown in Figure 5b and Figure 5c, respectively. Distribution of *P*(χ) and the structural details for *TSE*2 are shown in Figures 5d and 5e, respectively.

**Figure 5:**
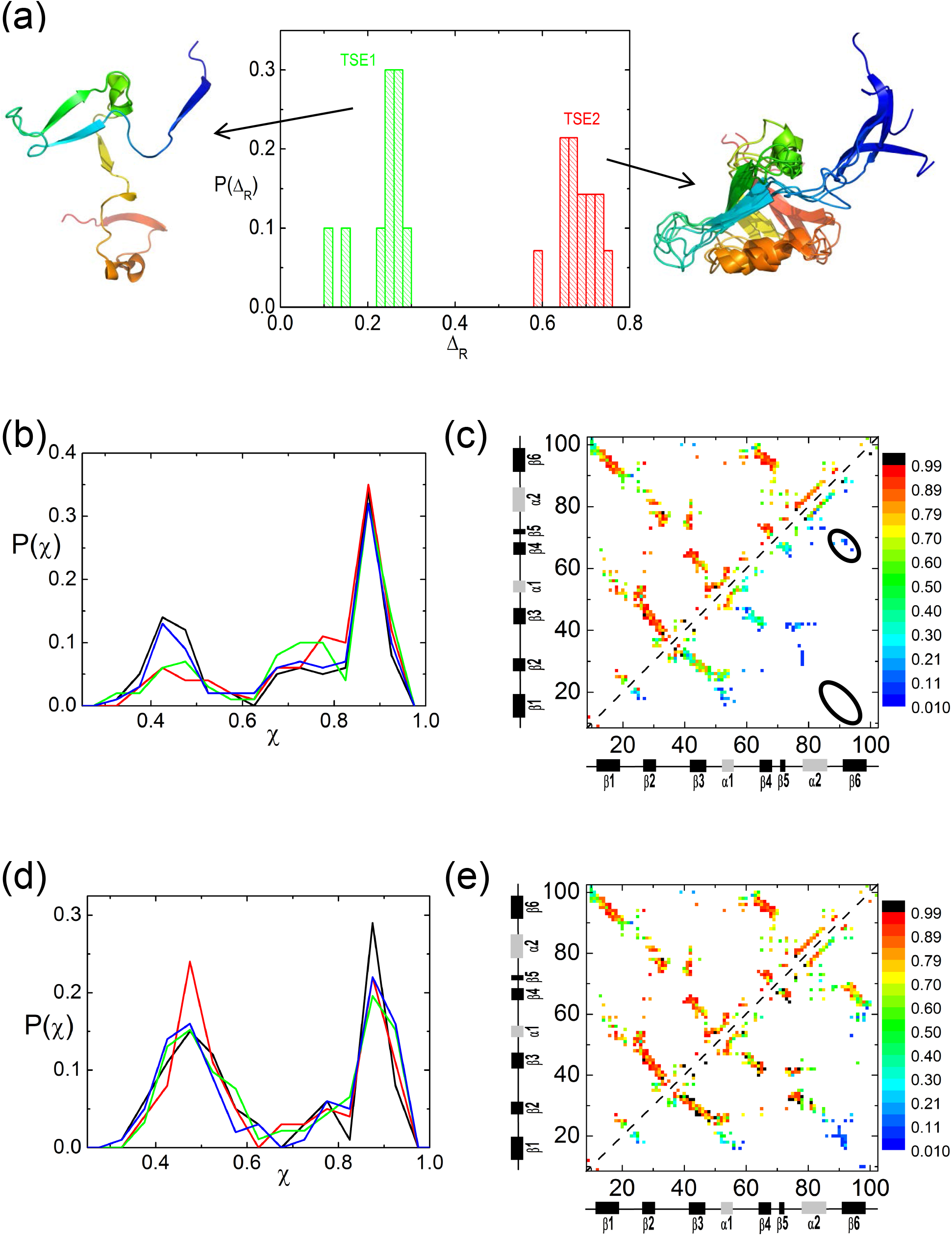
Quantifying the transition state ensembles. (a) Distribution *P*(Δ*_R_*), of the Δ_*R*_ = (Δ_*U*_ — *Δ_TSE_*)/(Δ_*U*_ – Δ_*N*_), which is the fraction of buried solvent accessible surface area relative to the unfolded structures. The average < Δ_*R*_ >= 0.23 for TSE1 and < Δ_*R*_ >= 0.68 for TSE2. These values coincide qualitatively with Tanford *β* parameters extracted from the observed Chevron plot. A few of the TSE1 (TSE2) structures are displayed on the left (right) respectively. (b)The distributions of χ computed from 400 simulation trajectories spawned from the transition state structures in TSE1. Data are shown for the four different structures. The distribution shows that roughly half of these trajectories go to the folded basin of attraction(P*_fold_* ≈ 0.5).(c) Contact map of the native state ensemble (upper left) and the one for the TSE1(lower right). The scale on the right gives the probability of contact formation. (d) and (e) Same as (b) and (c), respectively except the results are for TSE2.

The characteristics of the TSEs are experimentally described using the Tanford *β* parameter. By fitting the measured Chevron plots as linear function of [*C*] it has been shown that 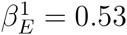 for TSE1, and 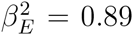 for TSE2^27^ and 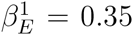 for TSE1 and 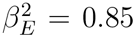 for TSE2^31^. It is generally assumed that *β* is related to the buried solvent accessible surface area (SASA) in the transition state. For the *TSE*s obtained in our simulations, we calculated the distribution *P(Δ_R_)* (see Figure 5a), where Δ_R_ = *(Δ_U_—ΔTSE)/(Δ_U_—ΔN)* with *Δ_U_, Δ_TSE_*, *Δ_N_* being the SASAs in the DSE ([*C*] = 8.0M), TSE, and the NBA ([*C*] = 0.0M), respectively. We found that the average 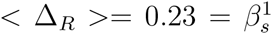 for *TSE*1 and 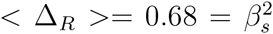 for *TSE*2, which qualitatively agree with the values inferred from experiments. The small value of *β*_1_ suggests that the *TSE*1 structures are more UBA-like whereas the *TSE*2 structures resemble the native state.

The structural details are revealed in Figure 5c, which shows the contact map for *TSE*1. It is clear that relative to the native state (top half in Figure 5c) the extent to which the structure is ordered in TSE 1 is modest (green) to low (blue). The contacts between *β* strands 1 — 6 — 4 have low formation probabilities as indicated by the two black circled regions. The secondary structures, *β*_23_ and two helices, are ordered to a greater extent. A representative structure for *TSE*1 displayed on the left of Figure 5a shows that the structure is expanded. Only *β*_23_ and the two helices have relatively high formation probability.

The contact map for *TSE*2 (Figure 5e) shows that the formation probabilities of contacts even between residues that are distant in sequence are high, which results in the ensemble of TSE2 structures being compact. The major blue region in the contact map indicates that *β*_16_ is still largely unstructured. Four superimposed representative structures from *TSE*2 are shown on the right of Figure 5a. The structures are native-like except that *β* strand 1 is not as well packed as in the native state. The lack of stabilizing interactions in *β*_16_ found in our simulations disagrees with the inferences from the all atom molecular dynamics simulations using measured Φ values as constraints^29^ (see below for additional discussion).

**Folding and collapse kinetics:** We calculated the collapse and folding rates at zero urea concentration from the folding trajectories, which were generated from Brownian dynamics simulations^39^. From 93 folding trajectories, the fraction of unfolded molecules at time *t*, is computed using 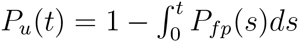, where *P_fp_(s)* is the distribution of first passage times. We fit *P_u_(t) = e^-tk_f_^* ( Figure 6a) with *k_f_* = 172s^-1^, a value that is about 60 times larger than found in the experiments (2.8^-1^)^27^. The kinetics of collapse of PDZ2 domain, shows that < *R_g_(t*) > decays with a single rate constant, *k_c_([C*]), the rate of collapse. The extracted values of *k_c_([C])* = 244s^-1^ from the data in Figure 6b is greater than kf ([C]), which shows compaction occurs ahead of folding.

**Figure 6:**
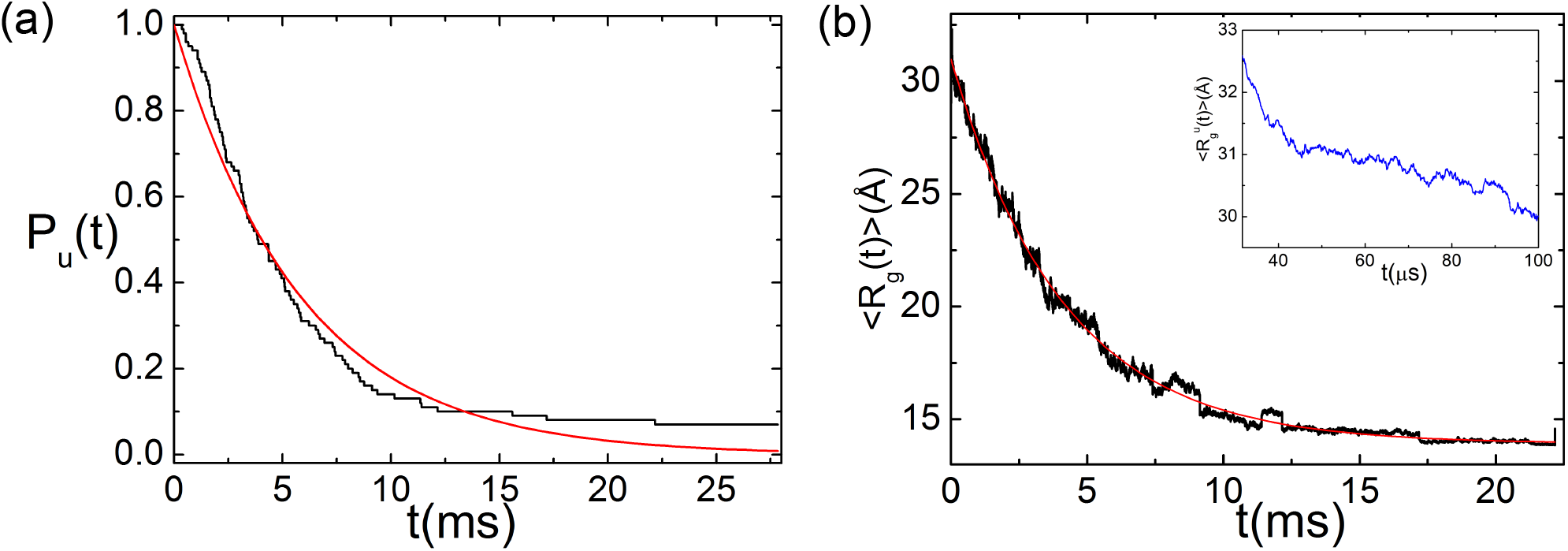
Folding and collapse kinetics. (a) Fraction of molecules that have not folded ([*C*] = 0.0*M*) as a function of time. The line is an exponential fit *(P_u_(t) = e^−^t/τ_F_* with τ_F_ = 5.8*ms*) to the data. (b) Collapse kinetics monitored by the time-dependent average < *R_g_ (t) >* as a function of t with the line giving an exponential fit to the data. The collapse time of PDZ2 is *τ_C_ = 4.1ms.*

**Thermodynamic and kinetic intermediates:** Analyses of the folding trajectories show that the folded state is reached through a kinetic intermediate state, *I*_1_ (Figures 7a, 7b, 7c, and 7d). This intermediate state, while not directly detected in experiment, is likely responsible for the downward curvature observed in the unfolding arm of the chevon plots^27^. We also find the presence of another kinetic intermediate states (*I*_2_) in some of the trajectories. A quantitative analyses allows us to classify the occurrence of *I*_1_ and *I*_2_ in the 93 folding trajectories. In 49 trajectories both *I*_1_ and *I*_2_ are found. Two trajectories in which this occurs are illustrated in Figures 7a and 7b. In Figure 7a, PDZ2 samples *I*_1_ and *I*_2_ only once before reaching the native state whereas in Figure 7b, it samples *I*_1_ and *I*_2_ more than once. In the second class of trajectories (44 out of 93) PDZ2 samples only the *I*_1_ (see Figures 7c and 7d for examples). Structures of *I*_1_ and *I*_2_ are displayed in Figure 7e.

**Figure 7:**
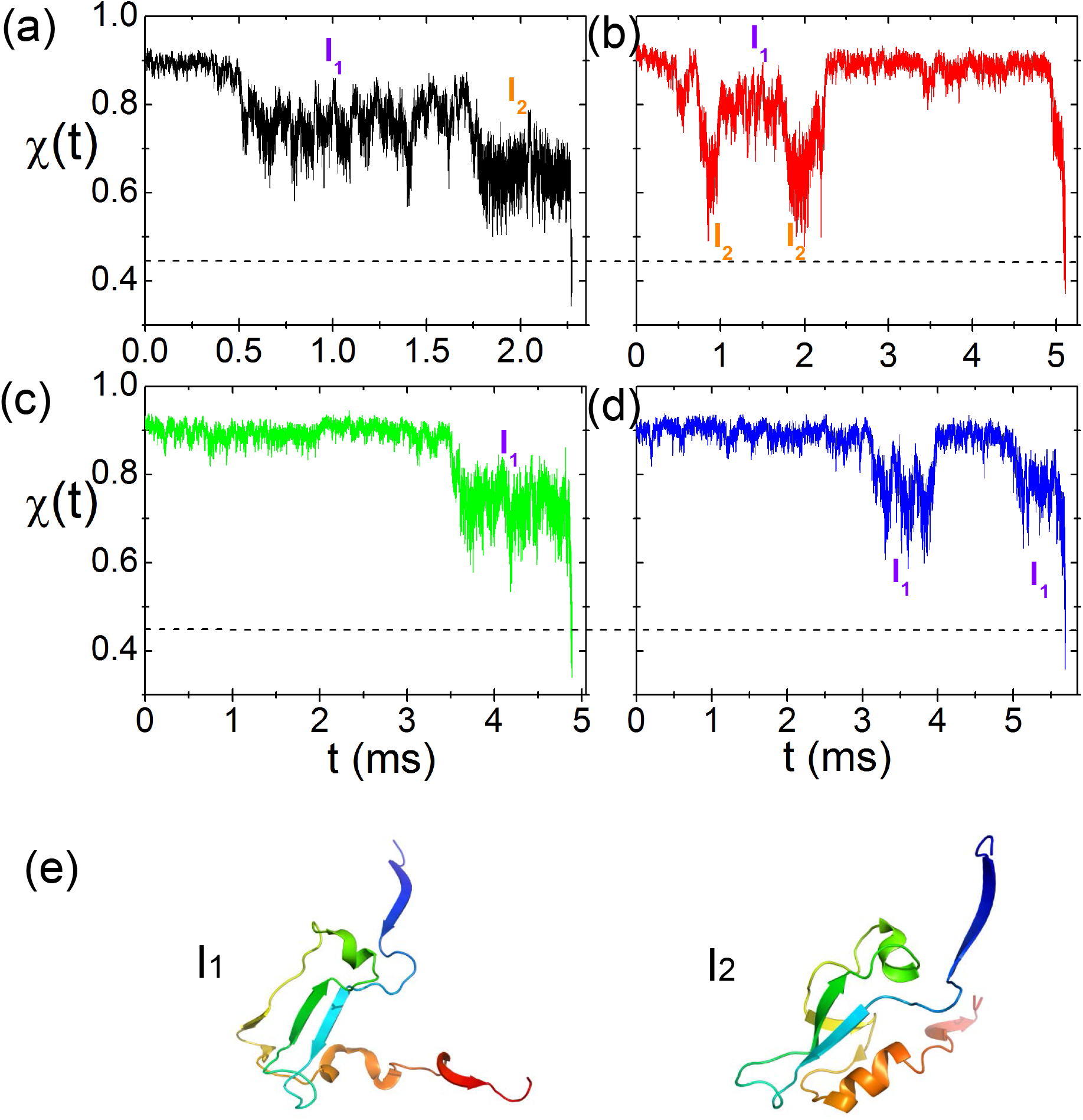
Two major folding pathways monitored by *χ* as a function of *t*. (a) and (b) show two representative trajectories showing that the native state is reached by sampling both *I_1_* and *I*_2_. (c) and (d) show two folding trajectories in which the polypeptide chain samples only *I*_1_, often multiple times, before reaching the folded state. (e) Structures for I1 and I2.

*I_EQ_* **and** *I*_1_ **are structurally similar:** To illustrate the structural similarity between *I_EQ_*, identified in the free energy profiles (Figures 1 and 4), and *I*_1_ in 100% of the kinetic folding trajectories, and *I*2 sampled in some of the folding trajectories, we computed the average fraction of native contacts formed by every residue, *f_Q_*, for the three states (Figures 8a and 8b). The correlation between *I_EQ_* and *I_1_,* shown in Figure 8c, is very high (R=0.995). Therefore, we surmise that *I*_1_ and *I_EQ_* are structurally identical. Quantitative results in Figures 8b and 8d and sample structures (Figure 8e) show that the structure of *I*_2_ differs from *I_EQ_*. Taken together these results show that both thermodynamic and kinetic intermediates can be sampled during the folding process although the former is far more prevalent.

**Figure 8:**
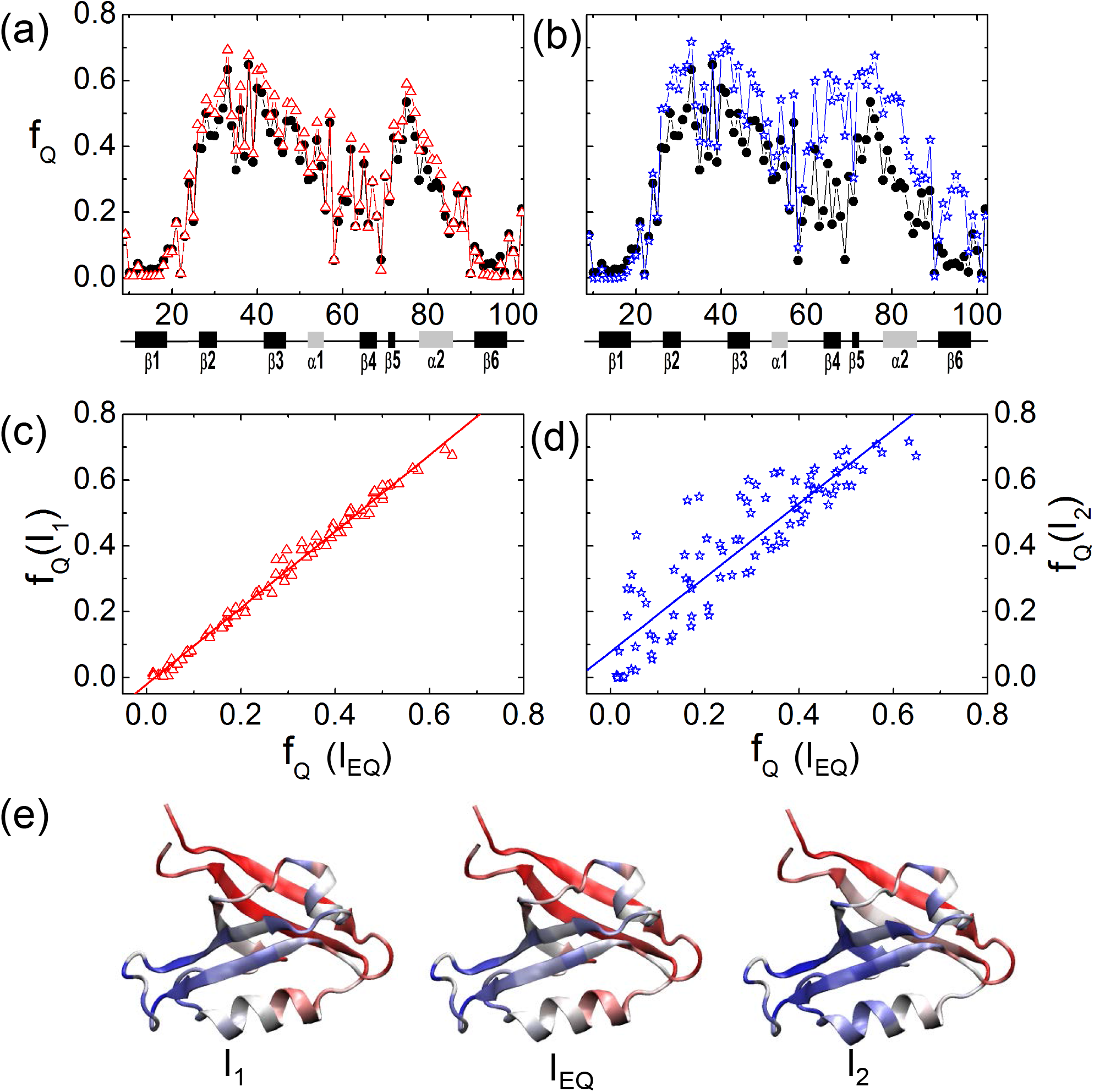
Comparing *I_EQ_*, *I*_1_, and *I*_2_. (a)Average fraction of native contacts formed for residues, *f_Q_*, for *I_EQ_* (black) and *I*_1_ (red). (b) *f_Q_* for *I_EQ_* (black) and *I*_2_ (blue). (c) Correlation between fQs for *I_EQ_* and *I*_1_. The correlation coefficient is near unity. (d) Relation between *f_Q_s* for *I_EQ_* and *I*_2_ with correlation coefficient ≈ 0.9. (e) Structures of the three intermediates.

**Figure 9:**
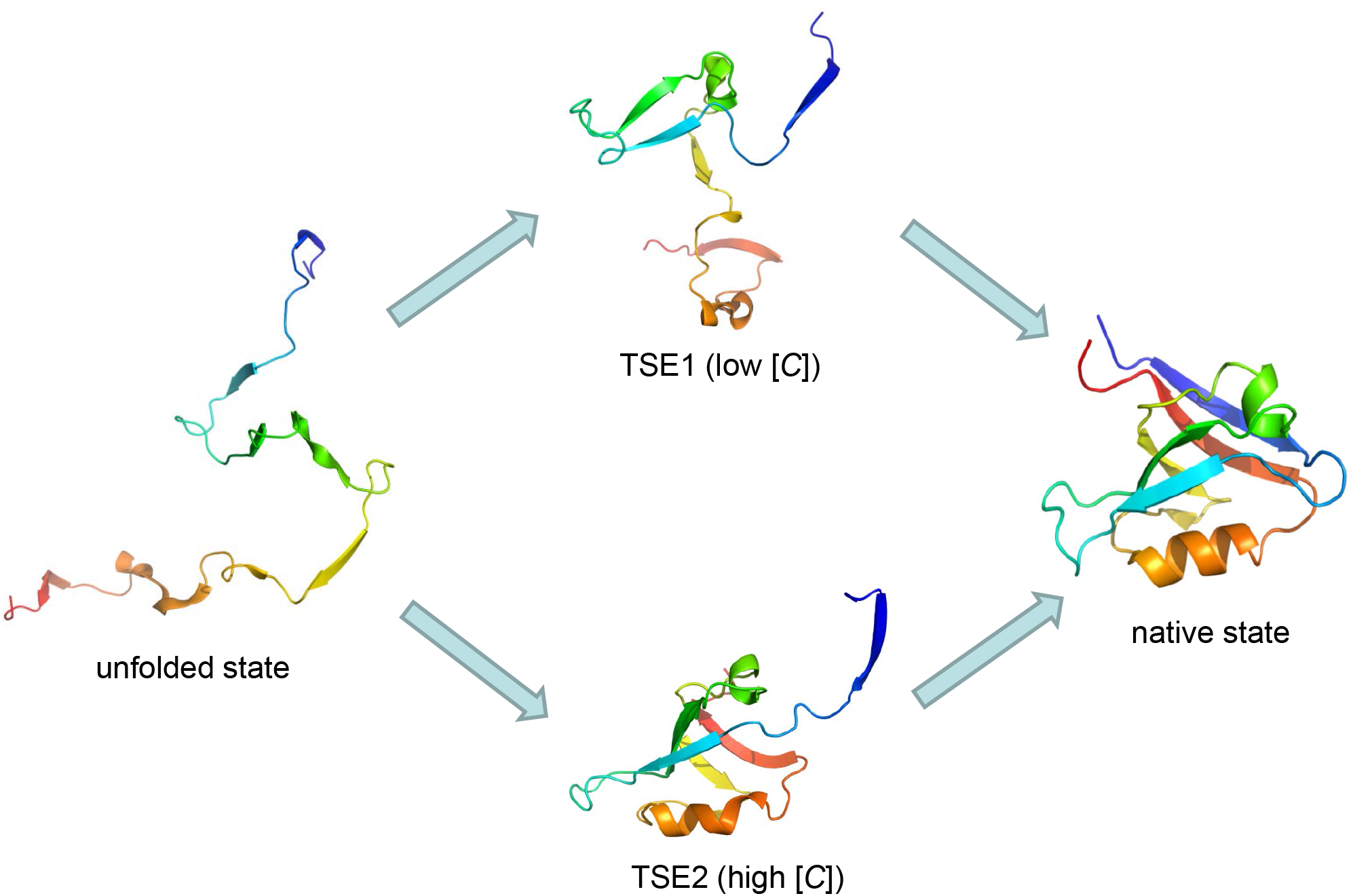
Table of contents.

## Discussion

**Minimum Energy Compact Structures (MECS):** The folding trajectories reveal that two major intermediates are sampled as PDZ2 folds. The equilibrium intermediate is found in all the trajectories whereas the kinetic intermediate is found in only a fraction of the trajectories. Two aspects of these findings, which are of general validity for folding, are worth pointing out. (i) Both *I*_1_ and *I*_2_ form on the time scale of collapse. In those trajectories in which *I*_2_ forms there frequent transitions between *I*_2_ and *I*_1_ (see Figure 7). In the process of making such transitions PDZ2 undergoes considerable expansion. (ii) The intermediates, *I_EQ_* and *I_1_*, are compact and contain native-like features. The major difference between *I*_1_ and the folded state is in the extent of structure in *β_1_, β_4_*, and *β_6_* as well as *α_2_.* The structure of *I*_1_ is similar to that found in *TSE*1, which follows from the Hammond postulate. These intermediates, which are like MECS (minimum energy compact structures)^49^ facilitate rapid folding. Although they are difficult to detect in ensemble experiments, single molecule pulling experiments using cycles of force increase and force quench can be used to detect MECS as has been done for Ubiquitin^50^. It would be interesting to do similar experiments on PDZ2 to directly observe *I*_1_ and *I*_2_ using force as a perturbation.

**Collapse and folding transition:** Is the size of the polypeptide chain of the unfolded state under folding conditions ([*C*] < *C_m_)* less than at [*C*] > *C_m_*? Theoretical arguments^46,51,52^, simulations^1^, and a number of single molecule FRET and SAXS experiments^19,53^ have answered the question in the affirmative whereas some SAXS experiments on protein L suggest that there is no evidence for polypeptide chain collapse, which is manifestly unphysical. The arguments in favor of collapse preceding folding is based on the following observations. The random coil state of a polypeptide chain with *N* residues is expected to be 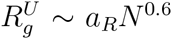 with *a_R_* ≈ 2.0Å. This estimate is likely to be an upper bound because even 8M urea is not a good solvent because even at these elevated concentrations the unfolded state has residual structure. As the denaturant concentration decreases the maximum decrease in *R_g_* is likely to have a lower bound *a_D_N*^0'5^. It cannot be maximally compact because if it were so then enthalpic interactions would drive these structures to the folded state for which *R_g_ « *a_N_N*^1/3^* with *a_N_* =3.0Å?. Thus, we surmise that 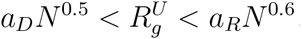. The reduction in *R_g_* of the unfolded state for *N* = 64 (protein L) is predicted to be between (5-10)% depending on the values of *a_R_* and *a_D_*. In PDZ2 (*N* = 94) we find that the unfolded state *R_g_* decreases by only 9% as [*C*] decreases (Figure 3a). Thus, due to finite *N* resulting in a small decrease in the unfolded state *R_g_* requires high precision SAXS measurements to measure changes in 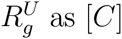 as [*C*] decreases. The errors in SAXS for protein L are far too large to accurately estimate 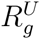, especially under native conditions.

The absence of detectable contracted form of the polypeptide chain in time resolved SAXS experiments protein L was used as further evidence that compact states are not formed in the folding of any single domain protein. Using theory^54-56^ it can be shown that collapse of the unfolded state occurs on time scales on the order of *τ_C_ ∼ τ_0_N^β^* where *β* is between 1.5 and 2.2, and 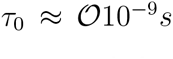. For PDZ2 we estimate that *τ_C_ ∼ 20μs*. Our simulations show that the contraction of the unfolded state occurs on time scale that does not exceed a maximum of ∼ 50*μs* (see the inset in Figure 6b). Theoretical estimate based on the scaling law above for collapse of the unfolded state of protein L (*N* = 64) a maximum of *τ_C_ ∼ 4μs.* Based on reconfiguration time measurements using Fluorescence Correlation Spectroscopy^57,58^ indicate that *τ_C_* for small proteins could be less by an order of magnitude compared to the theoretical estimate. A larger value of 20μs, much shorter than the folding time, for reconfiguration time has also been reported for protein L^59^. For the larger DHFR (*N* = 154) there is nearly a 23% reduction in 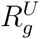 in ∼ 300μs^60^, which is also considerably shorter than the folding time. All the studies show that collapse of the unfolded state, which increases with *N*, is on the order of at most tens of *μs*. Thus, we conclude that the time resolution in the most recent SAXS experiments on protein L (4 ms)^61^, which is comparable to the folding time, is too long to shed light on compaction of the polypeptide chain. The presence of 15 His tags in the first study^62^ makes it difficult to ascertain the relevance for protein L. Indeed, the inability to accurately determine determine the characteristics of the unfolded state had also lead to erroneous conclusions about equilibrium collapse and kinetic foldability^63^, which has been corrected recently using smFRET experiments recently^64^.

Fast mixing experiments that simultaneously detect compaction and acquisition of structure on a number of proteins^65-67^ have produced ample evidence that collapse is an integral process of the folding process, as predicted by theory. Although it is likely that the simplified analysis of smFRET measurements overestimates the extent of collapse^68^, fast mixing SAXS experiments on a variety of proteins leave no doubt that the unfolded state is indeed more compact (albeit by only about ≈ 9%) under native conditions despite persistent claims to the contrary based on SAXS data based largely on one protein (protein L) with large errors. It is a pity that the erroneous conclusion has resulted in unnecessary obfuscation. What is needed are high precision data for single domain proteins spanning a range of *N* (say between 50-250), which cannot be easily obtained by SAXS alone^69^ but is more readily available in smFRET experiments.

**Detection of** *I_EQ_* **in PDZ2 in single molecule pulling experiments:** The downward curvature in the unfolding rate as a function [*C*] in PDZ2 implies that an intermediate is populated^27^. Single molecule pulling experiments are well suited to explore this finding more readily. Based on the free energy profile computed here and postulated elsewhere^31^, we suggest that unfolding by mechanical force (*f*) would give rise to downward curvature in a plot of *logk_u_(f*) versus *f*. At low forces the inner barrier separating the *NBA* and *I_EQ_* would dominate whereas at high forces the second outer barrier is relevant for mechanical unfolding. The two sequential barrier picture implies that there would be two transition state distances with a switch between the two occurring as *f* is increased. A plausible support for this argument comes from the observation that the Tanford *β*_T_ as a function of [*C*] exhibits a sigmoidal behavior (see Figure 4 in^31^) with *β_T_* changing from 0.35 at low [*C*] to 0.85 at high [*C*]. The scenario postulated here is distinct from that in SH3 domain in *[logk_u_(f),f]* plot exhibits an upward curvature, which is a signature of parallel unfolding pathways with a switch between the transition state ensembles^70^ as opposed to the predicted sequential barrier model for PDZ2. Laser optical tweezer experiments are ideally suited to test our prediction.

**Structures of the TSEs:**It is interesting to compare the structures of TSE1 and TSE2 obtained in this work with those reported earlier^29^. The structure of TSE1 (Figure 5) show interactions involving *β*_2_, *β*_3_, and formation of the two helices. This is in sharp contrast to the conclusion in^29^ suggesting that in TSE1 *β* strands 1 — 6 — 4 are structured whereas the other secondary structural elements are essentially disordered. Although the agreement between the predicted structures in our work and those reported in^29^ in TSE2 is better in that we find that it is native-like, there are crucial differences as well. In particular, the lack of stabilizing interactions involving *β*ι and *β*_6_ found in our simulations disagrees with the inferences drawn from the all atom molecular dynamics simulations using measured Φ values as constraints^29^. The differences are likely to be related to the completely different approaches used in the two studies. In^29^ the measured Φ values were used as constraints in standard all atom molecular dynamics simulations, which use inaccurate force fields. The procedure, while interesting, is not systematic in the sense the accumulation of errors both from the Φ values as well as the MD simulations is nearly impossible to quantify. More importantly, the putative TSE structures were not used to obtain the *P_fold_* values in the earlier study^29^, which casts doubt on whether the identified TSEs accurately represent the actual TSEs. We believe additional experiments, including perhaps double mutant cycles, would be needed to ascertain the nature of the TSEs in PDZ2.

## Concluding Remarks

We have used a phenomenological theory based on the MTM to simulate the folding of PDZ2 domain as function of temperature and urea. In addition to providing support to the folding mechanism discovered in experiments^27^ we have made a few testable predictions. (1) We have obtained a precise dependence of the melting point as a function of urea concentration, which can tested using standard calorimetry experiments. (2) The presence of high energy intermediate in the absence of added salt can be characterized using single molecule pulling experiments as shown using simulations for srcSH3 domain^71^ demonstrating that the excited state is sparsely *(O* ≈ – 5)% populated, which coincided with the findings based on NMR experiments^72^.

## Acknowledgements

We are grateful to Ben Schuler and Gilad Haran for their interest and useful comments. This work was supported in part by a grant from the National Science Foundation to DT through CHE 13-61946. Z.L. acknowledges partial financial support from the National Natural Science Foundation of China (11104015) and the Fundamental Research Funds for the Central Universities (2012LYB08).

